# Overproduction of Native and Click-able Colanic Acid Slime from Engineered *Escherichia coli*

**DOI:** 10.1101/2022.10.25.513687

**Authors:** Joanna C. Sadler, Richard C. Brewster, Alba F. González, Jessica S. Nirkko, Simon Varzandeh, Stephen Wallace

## Abstract

The fundamental biology and application of bacterial exopolysaccharides is gaining increasing attention. However, current synthetic biology efforts to produce the major component of *Escherichia sp*. slime, colanic acid, and functional derivatives thereof have been limited. Herein, we report the overproduction of colanic acid (up to 1.32 g/L) from D-glucose in an engineered strain of *E. coli* JM109. Furthermore, we report that chemically-synthesized L-fucose analogues containing an azide motif can be metabolically incorporated into the slime layer via a heterologous fucose salvage pathway from *Bacteroides sp*. and used in a click reaction to attach an organic cargo to the cell surface. This molecular engineered bio-polymer possesses enormous potential as a new tool for use in chemical, biological and materials research.

---

Exopolysaccharide slimes (EPS) are produced by many bacteria in response to external environmental stress. These polymeric carbohydrates are rapidly synthesized in the cell interior and exported to the cell surface to encapsulate the host in a protective slime-like layer. Colanic acid (CA) is the major exopolysaccharide produced by *Escherichia sp*. and has been shown to protect cells from toxic metal ions^1–3^, pathogenic microor-ganisms^[4]^, and, most recently, to delay the effect of aging in a microbe-associated host^4,5^. Colanic acid therefore has the potential to be used for a range of applications, including bio-material design, drug delivery and cosmetics research. In biocompatible chemistry, slime production has recently been shown to mitigate the damaging effects of hydrophobic metabolites and membrane-active surfactants. Nano-micelles containing a transition metal catalyst were shown to induce CA slime formation in *E. coli* NST74 and this increased cell viability^6^. Interestingly, slime production had no negative effect on (i) metabolite flux through an engineered styrene production pathway or (ii) the efficiency of a Fe-catalyzed cyclopropanation reaction at the cell surface. Slime formation could therefore be a viable approach to mitigate the otherwise toxic effects of engineered metabolites and/or non-enzymatic reactions to bacterial cells. Thus, the bio-production of CA and its derivatives from renewable resources via synthetic biology is an important challenge for industrial biotechnology.

The chemical structure of colanic acid is heterogeneous, yet contains a repeating unit of D-glucose (Glu), L-fucose (Fuc), D-galactose (Gal) and D-glucuronic acid (GluA), which together make the slime negatively charged (Figure 1A)^7–9^. The biosynthesis of CA is encoded by the 21-gene, 24 kb *wca* operon (formerly named *cps*) that is ubiquitous to *Escherichia sp*. (Figure 1B)^8^. Transcription is initiated via a JUMPStart-RfaH antitermination mechanism^10,11^ and is positively regulated by the transcriptional activators RcsA and RcsB^12,13^. Interestingly, plasmid-based overexpression of RcsA has been demonstrated as a viable engineering strategy to produce increased quantitates of CA from D-glucose (up to 350 mg/L)^1^.

**Fig. 1.**
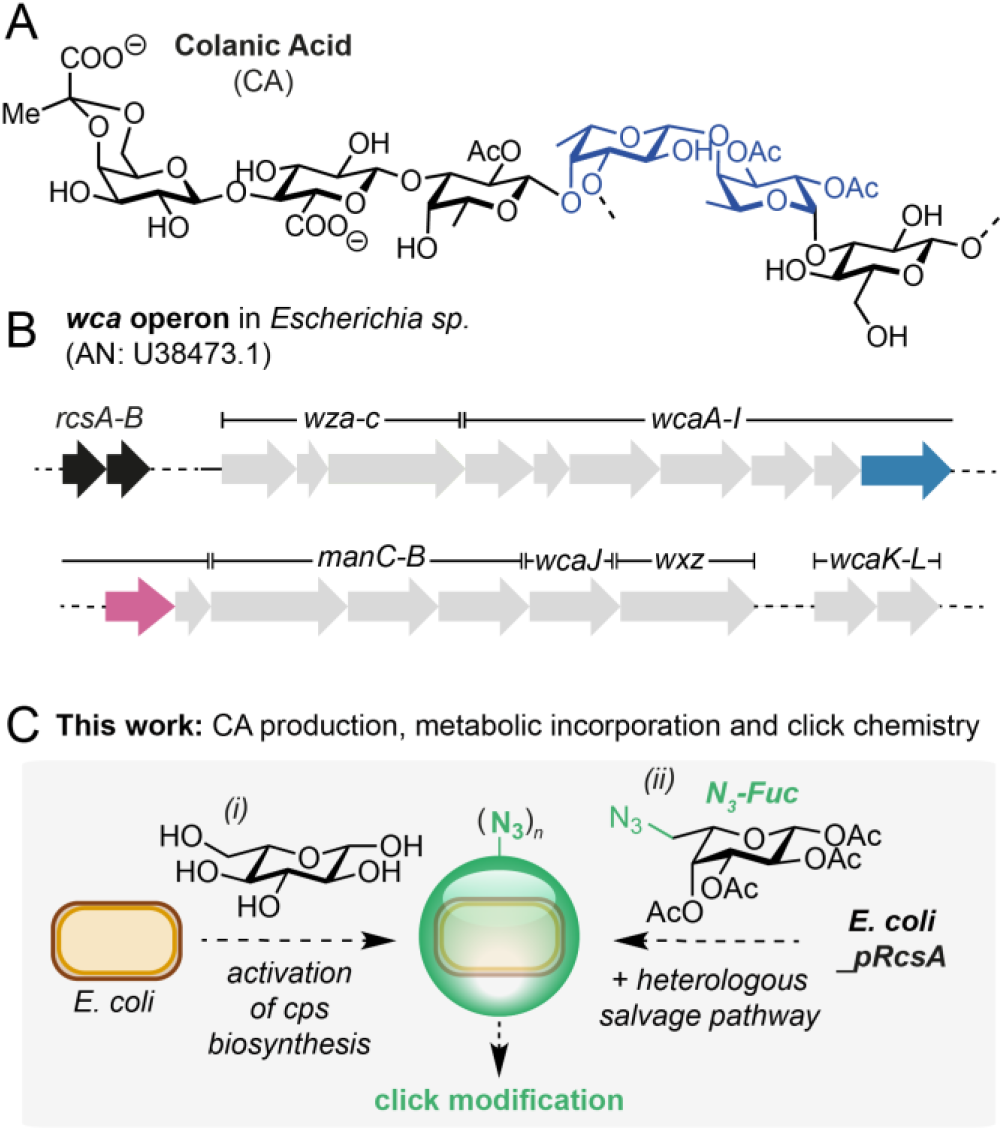
A) The structure of the major repeating unit in colanic acid (CA). L-fucose units are highlighted in blue. B) The wca operon in Escherichia sp. The positive regulators of colanic acid biosynthesis, rcsA/B, are highlighted in black, gmd and fcl are highlighted in blue and pink, respectively. C) Optimisation of CA production and incorporation of non-natural sugars. AN = accession number.

Inspired by these studies and recent interest in colanic acid from the biological community, we set out to increase the bio-production of CA and develop a method to modify its structure via the metabolic incorporation of an unnatural sugar. This is motivated by our interest in localizing non-enzymatic catalysts to the cell surface for use in new biocompatible reactions. Herein we report the high-level production of CA from D-glucose (1.3 g/L) in *E. coli* JM109Δ*waaF*_pRcsA under optimized fermentation conditions. Finally, we demonstrate the metabolic incorporation of the azide-containing unnatural sugar, GkiAZ, into CA using a fucose salvage pathway from *Bacteroides sp*.^14^ and conduct preliminary investigations into the use of chemically-engineered slime to localize cargo to the bacterial outer membrane via fluorescence detection.

We began by comparing RcsA-modified *E. coli* to other Gram-negative microorganisms that are known to produce acidics lime. We chose the organisms *Azotobacter vinelandii, Zooglea ramigera* and *Sinorrhizobium meliloti*, which are known to produce alginate, zooglan and succinoglycan polysaccharides, respectively^15–17^. *E. coli* BL21(DE3) and TOP10 cells transformed with a plasmid encoding RcsA (pRcsA) were used alongside an unmodified BL21(DE3) control. Slime formation was assayed by observing colony phenotype on agar plates supplemented with isopropyl β-D-1-thiogalactopyranoside (IPTG) to induce expression of RcsA (Figure 2A). Out of the *E. coli* strains tested, JM109_pRcsA was the only strain to produce a lustrous, shiny phenotype, which was indicative of CA overproduction. Conversely, BL21(DE3)_pRcsA and TOP10_pRcsA phenotypes were comparable to that of the JM109 empty vector negative control (JM109_pEdinbrick). This result was confirmed by measuring total carbohydrate and CA production in liquid cultures of BL21(DE3), TOP10 and JM109 transformed with pEdinbrick (empty vector control) or pRcsA (Figure 2B). Strain JM109_pRcsA gave >5-fold higher total carbohydrate titers and was the only strain to generate detectable levels of CA. This result was in agreement with the hypothesis that JM109 is deficient in Lon protease, which degrades RcsA^12,18^ and, unlike other common laboratory *E. coli* strains such as BL21(DE3) and Top10, does not carry *gal* mutations resulting in lower levels of CA intermediate UDP-D-galactose^19,20^.

**Fig. 2.**
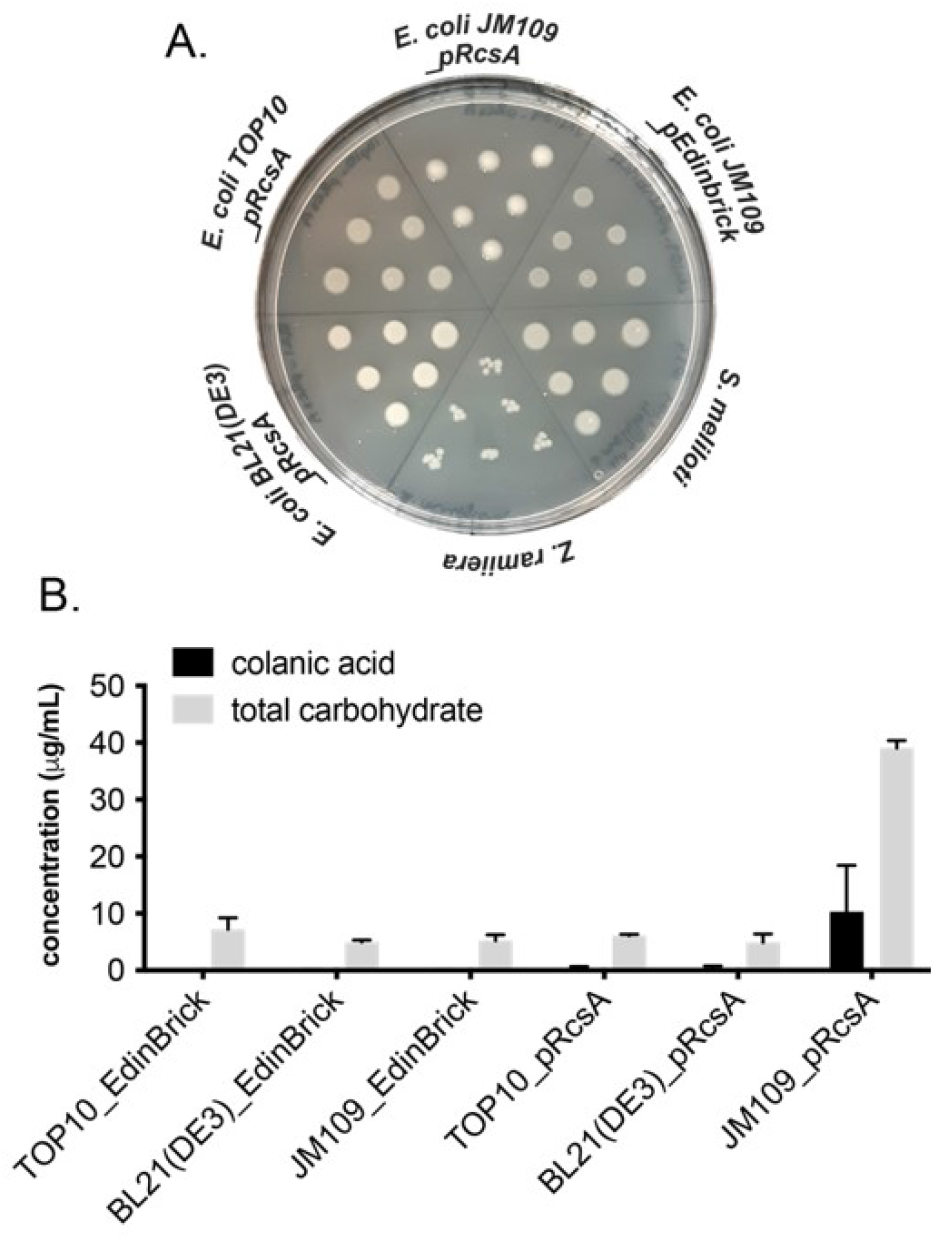
A) Solid phase screening for slime production on agar plates. Slime formation results in a lustrous, shiny colony phenotype. B) Liquid phase screening for total carbohydrate and colanic acid production by modified E. coli strains. Colanic acid was quantified by derivatization of extracted EPS with Cys.HCl and measuring the difference in absorbance at 427 nm and 396 nm. Total carbohydrate was quantified via the anthrone assay and measuring the absorbance at 620 nm. Data from quantitative experiments are presented as averages ofthree independent experiments to one standard deviation.

To maximize CA production by JM109_pRcsA, a series of optimization experiments was carried out. First, a media screen identified M9 and MDM, both minimal media, to provide the highest CA yields. Due to very slow growth rates and poor reproducibility in MDM, M9 was selected for all further investigations (Figure S6). Glucose concentration could be decreased from 5% w/v to 0.5% w/v without a significant effect on CA yield (Figure S7). The nitrogen source also had a profound effect on CA yields, with proline proving superior to both ammonium chloride and ammonium sulfate (Figures S8 and S9). The effect of the incubation temperature both in the presence and absence of trace amounts of Cu^2+^ and Fe^2+^ was also studied. A lower post-induction incubation temperature of 19 °C compared to 37 °C was found to increase CA yields to approximately 80% of the total carbohydrate content of the EPS (Figure 3A). The addition of Fe^2+^ and Cu^2+^ alone did not have a significant effect on CA yields at any temperature, however, they did increase growth rates of cultures. CA production over time was also studied. Both total carbohydrate and CA yields increased throughout the exponential growth phase, after which there was a small decrease in CA yields, despite a continued increase in total carbohydrate content (Figure 3B). Taking all of the optimization experiments together, the optimal conditions for CA production were concluded to be 90 hours incubation at 19 °C in M9 minimal media containing trace levels of Fe^2+^ and Cu^2+^, proline as the nitrogen source and 0.5% w/v glucose, yielding 419 mg/L CA (77 mg/L in unoptimized *E. coli* JM109). As a final optimization to further improve CA levels we knocked-out the *waaF* gene using a λ-Red recombinase. WaaF (also annotated as RfaF) is involved in carbohydrate tailoring during LPS biosynthesis in Gram-negative bacteria and Δ*waaF* strains have been shown to produce increased levels of EPS^[19]^. Indeed, this increased CA titers in the host strain to 719 mg/L, which increased further to 1.32 g/L when combined with *rcsA* overexpression (Figure 3C).

**Fig. 3.**
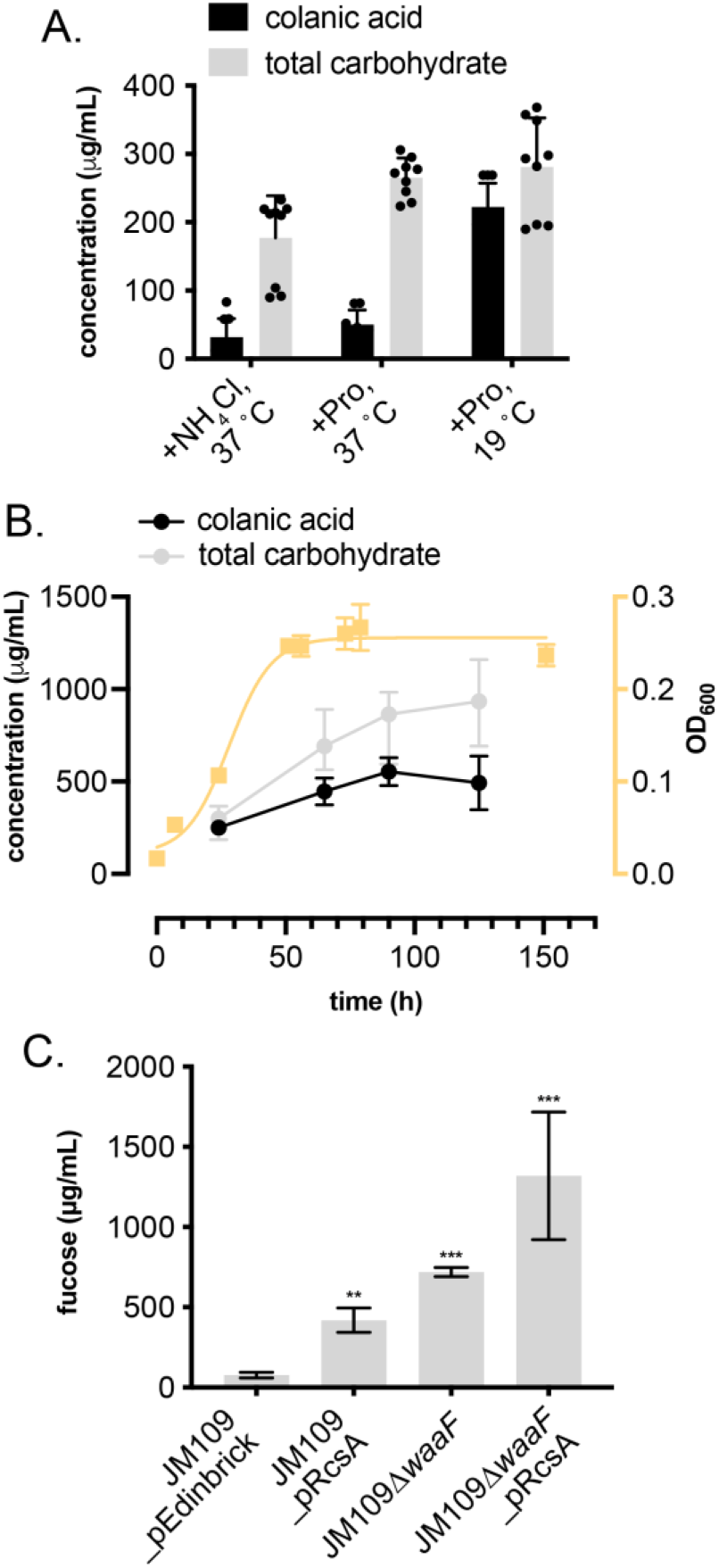
A) Temperature andN-source screen to increase total carbohydrate and CA production in E. coli JM109__pRcsA. B) CA production over time related to culture growth phase and total carbohydrate production. C) Strain optimization to enhance CA production. Fucose was quantified by acid hydrolysis and the L-Cys-phenol assay by spectrophotometric detection at 427 nm and 396 nm. Error bars represent the standard deviation of values from three independent experiments; *P<0.05 **P<0.005 ***p<0.005.

With conditions for increased levels of CA containing slime production in hand, we set out to functionalize the slime by metabolic incorporation of non-natural sugars. Whilst this is well established in mammalian cell lines,^[20]^ production of non-natural EPS analogues by *E. coli* is less established. A notable development in this field was the remodeling of bacterial polysaccharides by use of an exogenous sugar nucleotide salvage pathway^14^. In this work, the *de novo* GDP fucose pathway comprising GDP-mannose dehydratase (Gmd) and GDP-fucose synthetase (Fcl) was deactivated in a Δ*gmd-fcl* strain of *E. coli* (Figure 1B) and a salvage pathway activated by heterologous expression of the *fkp* gene, which encodes a bifunctional protein from *Bacteroides fragilis*^21^ with (i) fucokinase and (ii) GDP-fucose pyrophosphorylase activity. Following a similar strategy, we envisioned that unnatural azide-containing sugars could be introduced into bacterial slime via metabolic incorporation and that the resulting EPS could be functionalized using click chemistry. To this end, we prepared the fucose azide analogue Fuc-N_3_ in 5-steps from L-galactose via nucleophilic addition of sodium azide to the triflate generated from the primary alcohol of the corresponding *bis*-isopropylidene acetal, followed by deprotection in 30% overall yield (Figure 4). To determine whether the salvage pathway was required for the incorporation of Fuc-N_3_ we quantified fucose levels from Δ*gmd-fcl* strains grown in the presence and absence of fucose and Fuc-N_3_. The knockout strain was prepared using a λ-Red recombinase and then transformed with pRcsA and a plasmid encoding the *fkp* gene from *B. subtilis* (pFkp; Table S2). No fucose incorporation was observed in Δ*gmd-fcl* cells expressing *rcsA* when grown in the presence of glucose, glucose and fucose, or glucose and Fuc-N_3_, indicating the need for an alternative supply of GDP-fucose for CA biosynthesis in this strain. However, Δ*gmd-fcl* cells co-transformed with pRcsA and pFkp produced >100 mg/L CA when grown in the presence of Fuc but no CA was detected in the presence of glucose and Fuc-N_3_. This indicated that the salvage pathway from *Bacteroides sp*. was either unable to accept Fuc-N_3_ or was being poorly expressed in *E. coli* JM109. Analysis by SDS-PAGE indicated low levels of Fkp in cells and therefore we carried out a series of optimization experiments to increase protein expression. A screen of pre- and post-induction conditions revealed that Fkp levels could be significantly increased if cells were grown in LB media at 37 °C and then cooled to room temperature after induction with IPTG (Figure S10). Pleasingly, growth of *JM109Δgmd-fcl cells* expressing pRcsA under these optimized conditions and in the presence of glucose and Fuc-N_3_ resulted in the production of full length colanic acid EPS (Figure 5B). To confirm the incorporation of Fuc-N_3_ into the EPS we conducted a labeling experiment using a copper-catalyzed azide-alkyne cycloaddition (CuAAc) reaction (Figure 5A). The water-soluble ligand THPTA (*tris*((1-hydroxy-propyl-*1H*-1,2,3-triazol-4-yl)methyl)amine) was chosen instead of TBTA (*tris*(benzyltriazol-ylmethyl)amine) to increase penetration of the EPS, stabilize Cu(I) and disfavor oxidative side-reactions. The fluorescent al-kyne-containing dye 5-fluorescein-alkyne (5-FAM-alkyne) was chosen due to its known reactivity in the CuAAc reaction under aqueous conditions and visible excitation and emissions wavelengths at 485 nm and 520 nm, respectively. Therefore, azide EPS from JM109Δ*gmd-fcl*_pRcsA_pFkp cells was incubated with 5-FAM-alkyne in the presence of Cu(II)THPTA for 16 h, dialyzed and then analyzed by florescence spectroscopy. Pleasingly, fluorescent triazole-linked EPS was only detected in samples expressing *fkp* that had been incubated with Fuc-N_3_ and Cu(II) catalyst, confirming both the metabolic incorporation of Fuc-N_3_ through *wca* biosynthesis and the functional activity of the resulting EPS slime towards covalent attachment of an external small molecule (Figure 5C). Given the importance of EPS layers in bacterial biofilms, their pathogenicity in humans and the growing interest in colanic acid as a biocompatible polymeric material, this technology paves the way for the chemical design of new functional biomaterials for a variety of applications through metabolic incorporation and *in vivo* click chemistry in engineered bacteria. Colanic acid M-antigen is also tightly associated to the outer membrane after export by the Wzx flip-pase and therefore click-modified exopolysaccharide slimes could also provide a targeting methodology for the selective delivery of small molecules to bacterial cells in multicellular en-vironments.

**Fig. 4.**
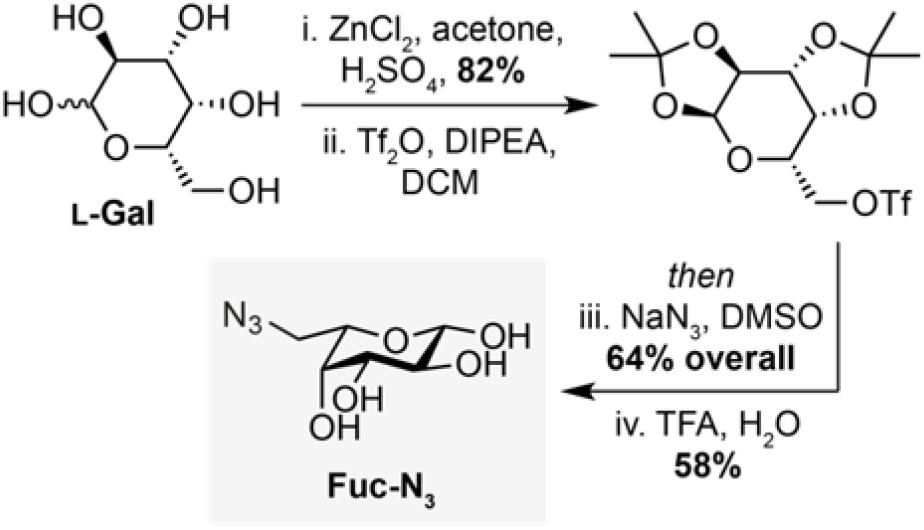
Synthesis of azide-containing fucose analog Fuc-N_3_.

**Fig. 5.**
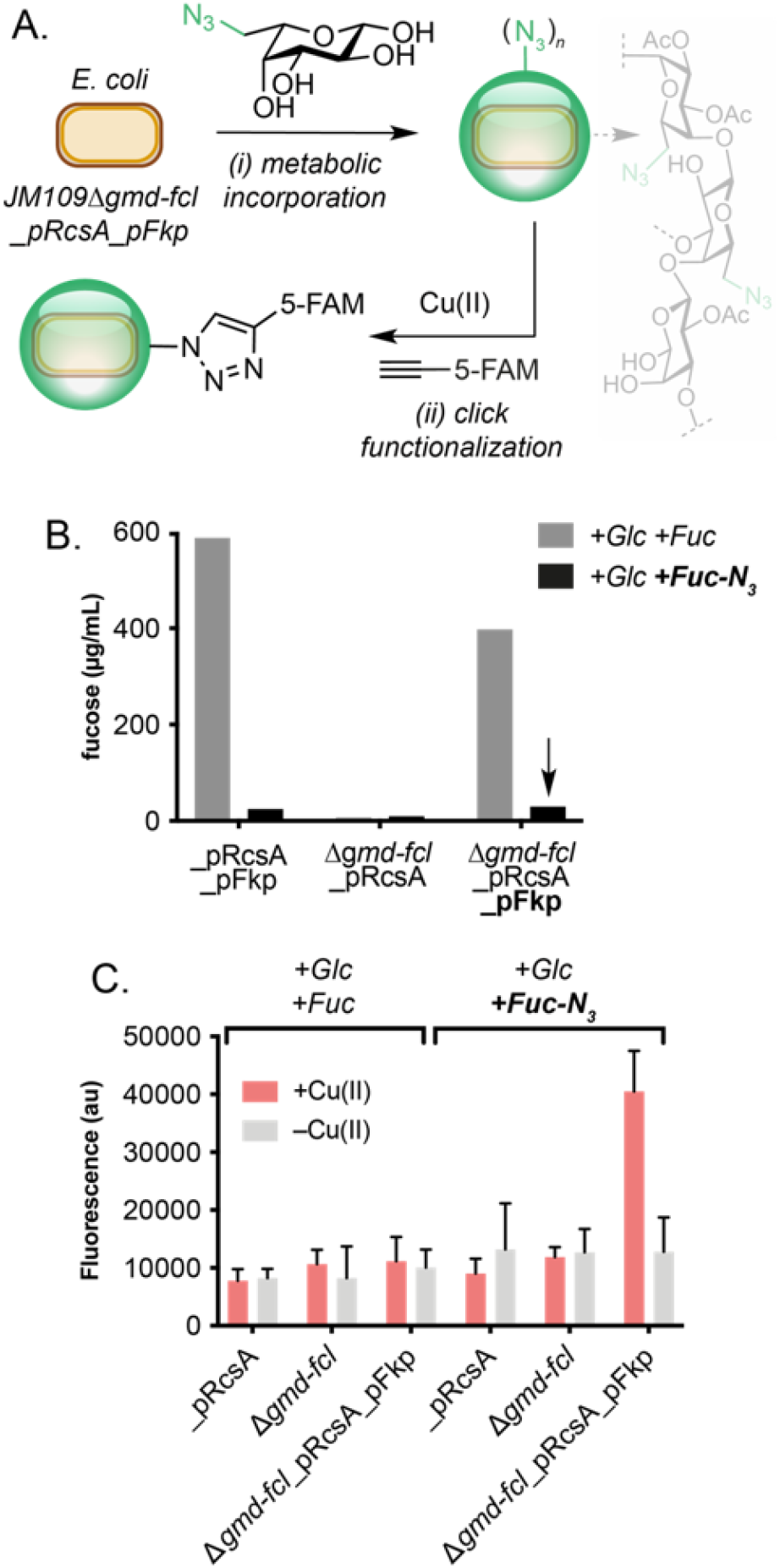
Click functionalization of azide-modified slime. A) Metabolic incorporation of Fuc-N_3_ in engineered E. coli JM109 and CuAAc labelling of a fluorescent dye. B) EPS production in engineered strains containing native and heterologous fucose salvage pathways when cultured in the presence of Fuc and Fuc-N_3_. C) Fluorescent read-out from click-modified EPS generated by strains engineered to metabolically incorporate Fuc-N_3_. Fucose was quantified by acid hydrolysis and addition of L-Cys, followed by spectrophotometric detection at 42 7 nm and 396 nm. Click reactions were performed using CuSO_4_ (100 μM), THPTA (500μM), 5-FAM-alkyne (50 μM), sodium ascorbate (5 mM) in potassium phosphate buffer (pH7, 100 mM), in 500 μL reaction volumes at 30 °C and 1000 rpm. Fluorescence was measured in a plate reader at λ_ex_=485 nm and λ_em_= 520 nm. THPTA = tris((1-hydroxy-propyl-1H-1,2,3-triazol-4-yl)methyl)amine. FAM = carboxyfluorescein. Data is presented as an average of three independent ex-perim ents to one standard deviation.

To conclude, this study reports the overproduction and chemo-enzymatic synthesis of modified bacterial exopolysaccharide slime from engineered *Escherichia coli*. High level production of colanic acid EPS was achieved through deregulation of the *wca* operon and redirection of carbohydrate away from LPS biosynthesis in *E. coli* JM109, and optimization of growth conditions (temperature and N-source). Unnatural azide-containing fucose (Fuc-N_3_) was synthesized and metabolically incorporated into the slime layer by deletion of native fucose pathways (Δ*gmd-fcl*) combined with heterologous expression of a fucose salvage pathway from *Bacteroides sp.*. When combined in a *JM109Δgmd-fcl_pRcsA_pFkp* engineered strain, azide modified EPS slime could be produced at room temperature, isolated and modified via a Cu-catalyzed azide-alkyne click reaction to produce a fluorescent product. This is the first production of such quantities of functionalized EPS slime from a metabolically engineered bacterium. Further metabolic engineering of this system to improve the production of both native and azide-modified EPS is currently underway in our laboratory. Overall, we anticipate broad utility of these bio-produced and chemi-cally-functionalized microbial polymers across the chemical and biological sciences.

## ASSOCIATED CONTENT

### Supporting Information

The Supporting Information is available free of charge on the BioRxiv website.

## AUTHOR INFORMATION

### Author Contributions

The manuscript was written through contributions of all authors. All authors have given approval to the final version of the manuscript.

### Funding Sources

J.C.S acknowledges a Discovery Fellowship from BBSRC (BB/S010629/1) and S.W acknowledges a Future Leaders Fellowship from UKRI (MR/S033882/1).

## Notes

### Competing Interest Statement

The authors have declared no competing interest.

