## Supplementary Information for "Overproduction of Native and Click-able Colanic Acid Slime from Engineered *Escherichia coli*"

#### 1 General materials and methods

Unless otherwise stated, starting materials and reagents were obtained from commercial suppliers and were used without further purification. All water used experimentally was purified with a Suez Select purification system (18 MΩ.cm, 0.2 µM filter).

##### 1.1 NMR Spectroscopy

Proton nuclear magnetic resonance spectra ($^1H$ NMR) were recorded using a Bruker AVA400, AVA500, Pro500 or AVA600 NMR spectrometer at the specified frequency at 298 K. Proton chemical shifts are expressed in parts per million (ppm, δ scale) and are referenced to residual protium in the NMR solvent (DMSO-$d_6$ = 2.50 ppm). Carbon nuclear magnetic resonance spectra ($^{13}C$ NMR) were recorded using a Bruker AVA400, AVA500, Pro500 or AVA600 NMR spectrometer at the specified frequency at 298 K. Chemical shifts are quoted in parts per million (ppm, δ scale) and are referenced to the carbon resonances of the NMR solvent (DMSO-$d_6$ = 39.5 ppm). Coupling constants, *J*, are measured to the nearest 0.1 Hz and are presented as observed. Data is represented as: chemical shift, integration, multiplicity (s = singlet, d = doublet, t = triplet, q = quartet, dd = doublet of doublets, m = multiplet and/or multiple resonances), coupling constant (*J*) in Hertz. All NMR solvents were purchased from commercial suppliers.

##### 1.2 Media recipes

**Lysogeny Broth (LB) Medium**

Bacto-tryptone (10 g/L), yeast extract (5 g/L) and NaCl (10 g/L) were dissolved in Milli-Q $H_2O$. LB was autoclaved at 121 ˚C for 20 minutes, cooled and stored at room temperature. LB agar was prepared using the same recipe but with the addition of agar (15 g/L).

**2XTY (Tryptone yeast) medium**

Bacto-tryptone (16 g/L), yeast extract (10 g/L) and NaCl (5 g/L) were dissolved in Milli-Q $H_2O$. 2XTY was autoclaved at 121 ˚C for 20 minutes, cooled and stored at room temperature.

**M9 minimal medium**

A 5X M9 stock solution was prepared as follows: $Na_2HPO_4 \cdot 12H_2O$ (85.5 g/L), $KH_2PO_4$ (15 g/L), NaCl (2.5 g/L), $NH_4Cl$ (5 g/L) and L-proline (27 g/L) were dissolved in Milli-Q $H_2O$ and autoclaved at 121 ˚C for 20 minutes. To prepare the final solution, filter sterilised $MgSO_4$ (2 mL, 1 M stock solution), $CaCl_2$ (2 mL, 50 mM stock solution), thiamine hydrochloride (0.8 mL, 50 mg/mL stock solution), $FeSO_4$ (1 mL, 10 µg/mL stock solution), $CuSO_4$ (10 mL, 1 µg/mL stock solution) and autoclaved glucose (25 mL, 20% w/v stock solution) were added to 200 mL 5X M9 stock solution and the final volume was adjusted to 1 L with sterile Milli-Q $H_2O$.

**M9 CA minimal medium**

M9 minimal medium was prepared as described above, with the addition of 25 g/L casamino acids being added to the 5X M9 stock solution, for a final concentration of 5 g/L casamino acids.

**MDM medium**

A 4X MDM stock solution was prepared as follows: $Na_2HPO_4{\cdot}7H_2O$ (64 g/L), $KH_2PO_4$ (15 g/L), NaCl (2.5 g/L) and $NH_4Cl$ (5 g/L) was dissolved in Milli-Q $H_2O$ and autoclaved at 121 °C for 20 minutes then cooled to room temperature. The mixture was diluted to 1X and filter sterilized thiamine hydrochloride (0.25 mM final concentration) and sterile glucose solution (1.25% w/v final concentration) were added. The mixture was supplemented with filter sterilised $MgSO_4$ (2 mL, 1 M stock solution), $CaCl_2$ (2 mL, 50 mM stock solution), thiamine hydrochloride (0.8 mL, 50 mg/mL stock solution), $FeSO_4$ (1 mL, 10 µg/mL stock solution) and $CuSO_4$ (10 mL, 1 µg/mL stock solution) and made up to 1 L final volume with Milli-Q water to make 1XMDM medium.

**YESCA medium**

0.5 g/L yeast extract and 5 g/L casamino acids were dissolved in Milli-Q $H_2O$ and autoclaved at 121 °C for 20 minutes then cooled to room temperature.

#### 2 Molecular biology

The *fkp* gene (Table 1) was codon-optimised for *E. coli* BL21(DE3) and synthesised using GeneArt™ (Thermo Scientific). Oligonucleotide primers were synthesised by Integrated DNA Technologies. Recombinant plasmid DNA was purified with a Miniprep Kit (Qiagen). *E. coli* strain JM109_pRcsA was kindly provided by the laboratory of Prof. French (University of Edinburgh). pRSFDuet-1 was purchased from Novagen. All restriction enzymes were purchased from Thermo Fisher as FastDigest™ enzymes. All restriction enzyme digests were carried out at 37 °C using FastDigest™ Green buffer. All plasmids were sequenced by Sanger sequencing at Edinburgh Genomics (Edinburgh, UK).

OneTaq 2X premix (New England Biolabs) was used for all colony PCR reactions, which contained 0.5 µM forward and reverse primers and water to a final volume of 12.5 µL. Phusion High-Fidelity DNA Polymerase (New England Biolabs) was used for all other PCR reactions, which contained 0.5 µM forward and reverse primers, 200 µM each dNTP, 10-20 ng template DNA and 1X HF buffer. T4 DNA ligase (Thermo Scientific) was used for all ligation reactions.

Colony PCR reactions were performed using the following conditions: initial denaturation (95 °C, 10 minutes), 35 thermal cycles (20 seconds denaturation at 95 °C, annealing at 50-65 °C for 30 seconds, and extension at 68 °C for 60 s/kb), and final extension (68 °C for 10 minutes). All standard Phusion PCR reactions were performed using the following conditions: initial denaturation (98 °C, 30 seconds), 30 thermal cycles (15 seconds denaturation at 98 °C, annealing at 50-72 °C for 20 seconds, and extension at 72 °C for 30 s/kb), and final extension (72 °C for 7 minutes). For agarose gel electrophoresis, agarose (1% w/v) TAE gels containing a 1 kB GeneRuler ladder (Thermo Scientific) were run at 100 V for 40 minutes and visualised using SYBR SafeTM. For SDS-PAGE, 12-well 12% acrylamide Bis-Tris NuPAGE gels (Thermo Scientific) containing an unstained Precision plus standard ladder (BioRad) were used to analyse samples. Gels were run in 1X MES buffer (Novagen) at 50 V for 30 minutes followed by 150 V for 2 hours.

#### 2.1 General procedure for the preparation of knockout strains

*E. coli* JM109(DE3)*Δgmd-fcl* and JM109*ΔwaaF* were generated via the reported protocol by Court et. al[1] with the following modifications: (i) the linear CamR cassette was purified via gel extraction and the coding DNA sequence was amplified a second time via PCR, (ii) cells were washed with glycerol (10% v/v aqueous solution) when preparing electrocompetent cells and were electroporated at 2.5 kV (200 Ω, 25 µFd), (iii) cultures were grown at 37 °C for 5 h following electroporation. The pSIM27 plasmid was used for recombineering and was obtained from the Court Lab (National Institute of Health, MD, USA). The chloramphenicol resistance cassette was constructed using the primers detailed in Table S1 using an annealing temperature of 60 ˚C under standard PCR conditions. Knockout colonies were confirmed via colony PCR using primers *gmd-fcl*_A-D or *waaF*_A-D (Table S1) as appropriate using an annealing temperature of 60 ˚C under standard colony PCR conditions.

***Table S1.*** *Primers used for the preparation of JM109(DE3)Δgmd-fcl and JM109ΔwaaF knockout strains.*

| **Primer number** | **Primer name** | **Primer sequence (5' to 3')** |
|---|---|---|
| 1 | CamR_*gmd-fcl* forward | TGTGACGGAAGATCACTTCG |
| 2 | CamR_*gmd-fcl* reverse | ACCAGCAATAGACATAAGCG |
| *3* | *gmd-fcl*_A | CGCGACTGTTCTCGACAATAAAGTCG |
| *4* | *gmd-fcl*_B | CACGACGATTTCCGGCAGTTTCTAC |
| *5* | *gmd-fcl*_C | GTAGAAACTGCCGGAAATCGTCGTG |
| *6* | *gmd-fcl*_D | GCTGGCAACCGATGTCTTTGTTGC |
| 7 | CamR_*waaF* forward | ATGGTGCCGTCCATTATTATCGCGGATGCCGGAAGTT AACGAAGCTATTCACCAGCAATAGACATAAGCG |
| 8 | CamR_*waaF* forward | GATAACCCTCCGCAGCGTCACCTTTACGCACTTTGTGA TAGCCGGTAATCTGTGACGGAAGATCACTTCG |
| 9 | *waaF*_A | CAGGCAGATCTGACAAATCTGCGC |
| 10 | *waaF*_B | CACGACGATTTCCGGCAGTTTCT AC |
| 11 | *waaF*_C | GTAGAAACTGCCGGAAATCGTCGTG |
| 12 | *waaF*_D | CGTCATAGTTCTCTGCTTGTAGCGC |

Figure S1 and Figure S2 demonstrate construction of the two knockout strains. Primers 3, 6, 9 and 12 anneal upstream and downstream of the *gmd-fcl* and *waaF* target genes, respectively. Together they yield a 2356 bp or 1686 bp PCR amplicon in unmodified *E. coli* K12 cells for *gmd-fcl* or *waaF* respectively. Upon insertion of the CamR cassette at the target locus, an approximately 1300 bp

(Δgmd-fcl) or 1700 bp (Δ*waaF*) amplicon is produced. Primers 4, 5, 10 and 11 anneal to the CamR cassette, producing PCR products only when the CamR cassette has been inserted into the genome.

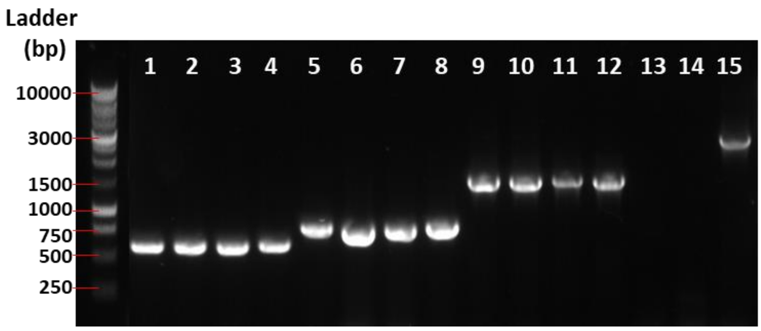

***Figure S1.*** *Analysis of colony PCR to confirm generation of E. coli JM109(DE3)Δgmd-fcl. Lanes 1-4: primers A+B, clones 1-4; Lanes 5-8: primers C+D, clones 1-4; Lanes 8-12: primers A+D, clones 1-4; Lane 13: A+B, JM109(DE3) control; Lane 14: C+D JM109(DE3) control; Land 15: A+D JM109(DE3)_pSIM27 control. Theoretical PCR product lengths: A/B: 578 bp C/D: 728 bp A/D: 1279 bp A/D (Control): 2356 bp.*

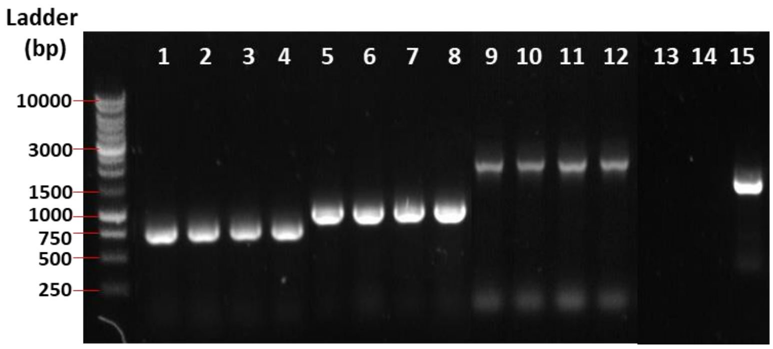

***Figure S2.*** *Analysis of colony PCR to confirm generation of E. coli JM109ΔwaaF. Lanes 1-4: primers A+B, clones 1-4; Lanes 5-8: primers C+D, clones 1-4; Lanes 8-12: primers A+D, clones 1-4; Lane 13: A+B, control Lane 14: C+D control Land 15: A+D JM109_pSIM27 control. Theoretical PCR product lengths: A/B: 768 bp; C/D: 943 bp; A/D: 1686 bp; A/D (Control): 1684 bp.*

#### 2.2 Strains and plasmids

***Table S2.*** *Table of plasmids used in this study*

| **Name** | **Description** | **Reference** |
|---|---|---|
| pEdinbrick | Control plasmid: empty pSB1A2-derived vector with AmpR and pUC19-derived pMB1 origin of replication. | [2] |
| pRcsA | EPS overproduction plasmid: pSB1A2 backbone containing the *lac* promotor (part Bba J33207 [from Design of BioBricks, http://syntheticbiology.org/BioBricks/Part fabrication.html)] and *E. coli* K12 *rcsA* gene. | [3] |
| pFkp | Fkp expression plasmid: pRSFDuet1 backbone with *fkp* gene inserted into multiple cloning site 1 using NcoI and NotI restriction sites. | This study |

| | |
|---|---|
| | *fkp* (from *Bacteroides fragilis*) gene sequence codon optimized for expression in *E. coli*:<br>ATGGCAATGCAGAAACTGCTGAGCCTGCCGAGCAATCTGGTTCAGAGCTTTCA<br>TGAACTGGAACGTGTTAATCGTACCGATTGGTTTTGTACCAGCGATCCGGTTG<br>GTAAAAAACTTGGTAGCGGTGGTGGCACCAGCTGGCTGCTGGAAGAATGTTA<br>TAATGAATACAGTGATGGTGCGACCTTTGGTGAATGGCTGGAAAAAGAAAAA<br>CGTATTCTGCTGCATGCCGGTGGTCAGAGCCGTCGTCTGCCTGGTTATGCACC<br>GAGCGGTAAAATTCTGACACCGGTTCCGGTTTTTCGTTGGGAACGTGGTCAGC<br>ATCTGGGTCAGAATCTGCTGTCACTGCAGCTGCCGCTGTATGAAAAAATCATG<br>AGCCTGGCACCGGATAAACTGCATACCCTGATTGCAAGCGGTGATGTTTATAT<br>TCGTAGTGAAAAACCGCTGCAGAGCATTCCGGAAGCAGATGTTGTTTGTTATG<br>GTCTGTGGGTTGATCCGAGCCTGGCGACCCATCATGGTGTTTTTGCAAGCGAT<br>CGTAAACATCCGGAACAGCTGGATTTTATGCTGCAGAAACCGAGTCTGGCAGA<br>ACTGGAAAGCCTGAGCAAAACCCACCTGTTTCTGATGGATATTGGTATTTGGC<br>TGCTGAGCGATCGTGCAGTTGAAATTCTGATGAAACGTAGCCATAAAGAAAGC<br>AGCGAAGAACTGAAATATTACGATCTGTATAGCGATTTTGGTCTGGCACTGGG<br>CACCCATCCGCGTATTGAAGATGAAGAAGTTAATACCCTGAGCGTTGCAATTC<br>TGCCGCTGCCTGGTGGCGAATTTTATCATTATGGTACAAGCAAAGAACTGATC<br>AGCAGCACCCTGAGTGTTCAGAATAAAGTTTATGATCAGCGTCGCATCATGCA<br>CCGTAAAGTTAAACCGAATCCGGCAATGTTTGTGCAGAATGCAGTTGTTCGTA<br>TTCCGCTGTGTGCAGAAAATGCAGATCTGTGGATTGAAAACAGCCATATTGGT<br>CCGAAATGGAAAATTGCAAGCCGTCATATTATCACCGGTGTTCCGGAAAATGA<br>TTGGAGCCTGGCCGTTCCGGCAGGCGTTTGTGTTGATGTTGTTCCGATGGGTG<br>ATAAAGGTTTTGTTGCACGTCCGTATGGTCTGGATGATGTTTTTAAAGGTGATC<br>TGCGTGATAGCAAAACCACACTGACCGGTATTCCGTTTGGCGAATGGATGAGC<br>AAACGTGGTCTGAGCTATACCGATCTGAAAGGCCGTACCGATGATCTGCAGGC<br>AGTTAGCGTTTTTCCGATGGTTAATAGCGTTGAAGAACTGGGTTTAGTTCTGCG<br>TTGGATGCTGAGTGAACCGGAACTGGAAGAAGGTAAAAACATTTGGTTACGC<br>AGCGAACATTTTAGCGCAGATGAAATTAGTGCCGGTGCAAATCTGAAACGTCT<br>GTATGCACAGCGTGAAGAATTTCGTAAAGGTAATTGGAAAGCACTGGCCGTG<br>AATCATGAAAAAAGCGTTTTTTATCAGCTGGATCTGGCAGATGCAGCCGAAGA<br>TTTTGTTCGTTTAGGTCTGGATATGCCGGAACTGCTGCCGGAAGATGCACTGC<br>AGATGAGCCGTATTCATAATCGTATGCTGCGTGCCCGTATTCTGAAACTGGAT<br>GGTAAAGATTATCGTCCGGAAGAACAGGCAGCATTTGATCTGCTGCGTGATG<br>GTCTGCTGGATGGCATTAGCAATCGTAAAAGCACCCCGAAACTGGACGTTTAT<br>AGCGATCAGATTGTTTGGGGTCGTAGTCCGGTTCGTATTGATATGGCAGGCGG<br>TTGGACCGATACACCGCCTTATAGCCTGTATAGTGGTGGTAATGTTGTTAACCT<br>GGCCATTGAACTGAATGGTCAGCCTCCGCTGCAGGTTTATGTTAAACCGTGTA<br>AAGATTTTCATATCGTCCTGCGCTCAATCGATATGGGTGCAATGGAAATTGTTA<br>GCACCTTTGATGAACTGCAGGACTACAAAAAAATCGGTAGCCCGTTTAGTATT<br>CCGAAAGCAGCACTGTCACTGGCAGGTTTTGCCCCTGCATTTAGTGCAGTTAG<br>CTATGCAAGCCTGGAAGAACAACTGAAAGATTTTGGTGCAGGTATTGAAGTTA<br>CCCTGCTGGCAGCAATTCCTGCAGGTAGCGGTCTGGGCACCAGTAGCATTCTG<br>GCAAGCACCGTTCTGGGTGCCATTAATGATTTTTGTGGTCTGGCGTGGGATAA<br>AAACGAAATTTGTCAGCGTACCCTGGTTCTGGAACAGTTACTGACCACAGGTG<br>GTGGTTGGCAGGATCAGTATGGTGGTGTTCTGCAGGGTGTTAAACTGCTGCA<br>GACCGAAGCCGGTTTTGCACAGAGTCCGCTGGTTCGTTGGCTGCCGGATCACC<br>TGTTTACCCATCCGGAATATAAAGATTGTCATCTGCTGTATTATACCGGCATTA<br>CCCGTACCGCAAAAGGTATTCTGGCCGAAATTGTGAGCAGCATGTTTCTGAAT<br>AGCAGCCTGCATCTGAACCTGCTGTCAGAAATGAAAGCACATGCACTGGATAT<br>GAATGAAGCAATTCAGCGTGGTAGCTTTGTTGAATTTGGTCGTCTGGTGGGTA |

| | |
|---|---|
| | AAACCTGGGAACAGAACAAAGCCCTGGATAGCGGCACCAATCCTCCGGCAGTTGAAGCCATTATTGATCTGATCAAAGATTATACCCTGGGCTATAAACTGCCAGGTGCCGGTGGCGGTGGTTATCTGTATATGGTTGCAAAAGATCCGCAGGCAGCAGTGCGTATTCGTAAAATCCTGACCGAAAATGCTCCGAATCCGCGTGCACGTTTTGTTGAAATGACCCTGTCAGATAAAGGCTTTCAGGTTAGCCGTAGCTAA |

#### 3 Quantification methods

##### 3.1 Colanic acid quantification in cuvettes

Quantification of colanic acid was carried out based on the protocol reported by from Obadia and co-workers[4] which measures fucose concentration, a constituent of EPS. Purified EPS samples were diluted in Milli-Q $H_2O$ depending on their predicted concentration to fit within the standard curve. Triplicate runs of the colonic acid assay were carried out by mixing 444 µL of the diluted samples with 2 mL of $H_2SO_4/H_2O$ (6:1 v/v) in a glass tube. The mixture was then heated to 95 °C for 30 mins and then cooled to room temperature. For each sample, 20 µL of (a) a freshly prepared cysteine hydrochloride (Cys·HCl, 3% (w/v) solution) of (b) Milli-Q water was added to a polystyrene cuvette. 1 mL of the sample was added and the absorbance of each cuvette was measured at both 396 nm and 427 nm. The absorbance measurements at both 396 and 427 nm without Cys·HCl were subtracted from those with Cys·HCl to provide background corrected $A_{396}$ and $A_{427}$ values. The final absorbance values were calculated by subtracting $A_{427}$ from $A_{396}$. The result was directly correlated to fucose concentration by using a fucose standard curve ranging from 5 µg/mL to 100 µg/mL.

##### 3.2 Colanic acid quantification using plate reader

The colanic acid quantification assay was successfully miniaturized for high throughput analysis using a plate reader. Purified EPS samples were diluted in Milli-Q $H_2O$ depending on their predicted concentration to fit within the standard curve. 111 µL of the diluted EPS mixture was added to 500 µL $H_2SO_4/H_2O$ (6:1 v/v) and the mixture was heated to 95 °C for 30 minutes then cooled to room temperature. For each sample, to a well of a flat-bottomed 96-well place was added (a) 5 µL cysteine hydrochloride (Cys·HCl, 3% stock solution) or (b) 5 µL Milli-Q water. To each of these was added 200 µL of the cooled acidified EPS mixture. The absorbance of both (a) and (b) was measured at 396 and 427 nm. The absorbance measurements at both 396 and 427 nm without Cys·HCl were subtracted from those with Cys·HCl to provide background corrected $A_{396}$ and $A_{427}$ values. The final absorbance values were calculated by subtracting $A_{427}$ from $A_{396}$. The result was directly correlated to fucose concentration by using a fucose standard curve ranging from 5 µg/mL to 100 µg/mL. A separate standard curve was prepared in parallel to the samples for each analytical run to account for differences in incubation times. A representative standard curve is shown in Figure S3.

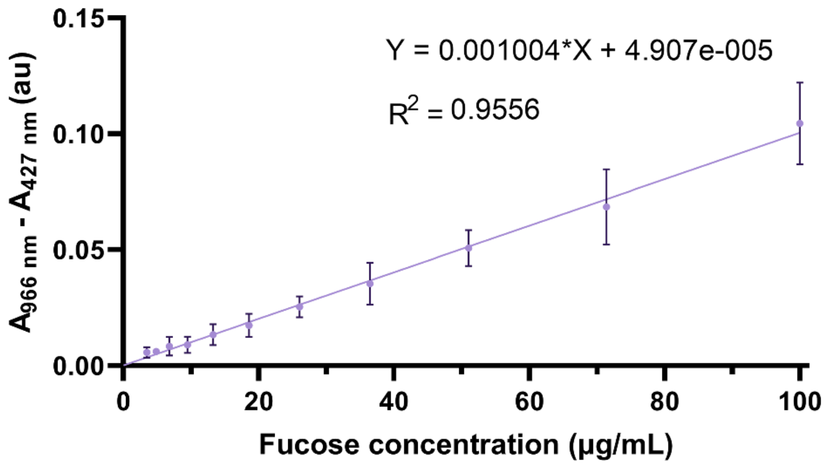

***Figure S3.*** *Representative standard curve for the quantification of fucose concentration.*

##### 3.3 Quantitative analysis of total carbohydrate content quantification

The total carbohydrate content of the samples was quantified by the anthrone-sulphuric acid assay based on a protocol from Rondel and coworkers[5]. The purified EPS samples were diluted in Milli-Q $H_2O$ depending on their expected concentration to fit within the standard curve, and 400 µL aliquots were added to a glass vial for each sample. To this was added 800 µL of a freshly prepared anthrone solution (2% w/v in 96% aq. $H_2SO_4$). The mixture was heated at 60 °C for 30 minutes then cooled to room temperature. The absorbance of the resulting solution at 620 nm was measured and correlated to glucose concentration using a standard curve ranging from 5 µg/mL to 100 µg/mL. A representative standard curve shown in Figure S4.

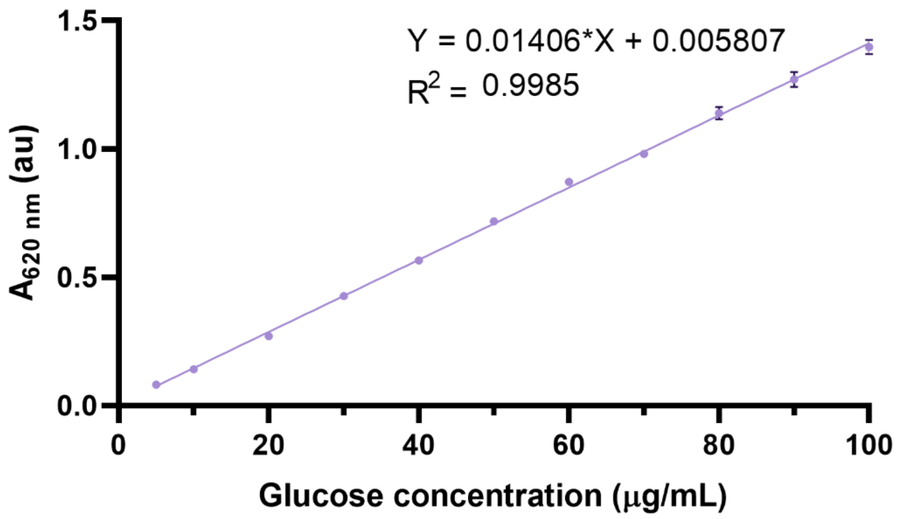

***Figure S4.*** *Representative standard curve for the quantification of glucose concentration.*

#### 4 EPS production

##### 4.1 Optimisation of colanic acid production

**Glucose concentration**

*E. coli* JM109_pRcsA was grown in minimal M9 or 1xMDM media supplemented with glucose at concentrations of 5% (w/v) or 0.5% (w/v). All cultures were incubated at 37 °C with rotary shaking at 220 rpm 24 hours post-induction.

**Nitrogen source**

M9 and 1xMDM media were prepared according to the recipes in Section 1.2, but excluding the nitrogen source. Ammonium chloride or ammonium sulphate were added at a concentration of 1 g/L and proline was added either at a concentration of 1 g/L or 5.4 g/L. All cultures were incubated at 37 °C and 220 rpm for 24 hours post-induction.

**Temperature and addition of trace metals**

M9 with either $NH_4Cl$ at 1 g/L or proline at 5.4 g/L was supplemented with $FeSO_4{\cdot}7H_2O$ 0.001 g/L and $CuSO_4{\cdot}5H_2O$ 0.001 g/L to test their effect on the growth and production of exopolysaccharides by *E. coli* JM109_pRcsA. The cultures were incubated either at 37 °C or 19 °C and 220 rpm until they reached an $OD_{600}$ = 0.6 at which point they were induced with isopropyl β-D-1- thiogalactopyranoside (IPTG) (0.5 mM final concentration). Further incubation took place in the same conditions over 24 hours, except for the cultures previously grown at 37 °C were either maintained at the same temperature or incubated at 19 °C.

**Time**

Time course experiments were carried out in M9 minimal media supplemented with proline at 5.4 g/L as the nitrogen source, trace metals $FeSO_4{\cdot}7H_2O$ and $CuSO_4{\cdot}5H_2O$ at a final concentration of 0.001 g/L and 5%w/v glucose as the carbon source. All cultures were incubated at 19 °C and 220 rpm for approximately 24 hours until the induction point. Further incubation was carried out under the same conditions for 65, 90 or 115 hours. Growth curves were obtained by taking 200 µL aliquots at different time points and measuring the $OD_{600}$.

##### 4.2 General procedure colanic acid production under optimized conditions

LB (10 mL) containing ampicillin (100 µg/mL) was inoculated with a single colony from freshly streaked plates of JM109_pRcsA or JM109_pEdinbrick and incubated overnight at 37 °C with shaking (220 rpm). The next day, M9 EPS media (2 or 5 mL depending upon experiment) containing ampicillin (100 µg/mL) was inoculated with 2% v/v overnight culture and incubated at 19 °C for 24 hours. IPTG was added (100 µM final concentration) and cultures were incubated at 19 °C for a further 72-96 hours (unless otherwise specified).

##### 4.3 General procedure for production of azide-labelled colanic acid

LB (10 mL) containing ampicillin (100 µg/mL) was inoculated with a single colony from freshly streaked plates of JM109(DE3)Δ*gmd-fcl*_pRcsA_pFkp and incubated overnight at 37 °C with shaking (220 rpm). The next day, M9 EPS media (2 or 5 mL) containing ampicillin (100 µg/mL) and 0.1%w/v Fuc-$N_3$ was inoculated with 2% v/v overnight culture and incubated at 37 °C for until $OD_{600}$=0.3-0.4 . IPTG was added (500 µM final concentration) and cultures were incubated at 19 °C for a further 72 hours.

##### 4.4 General procedure for fluorescent labelling of azide-labelled colanic acid

Cultures containing azide-labelled colanic acid were diluted to $OD_{600}$=1 in sterile phosphate buffered saline (PBS) and dialysed against PBS for 24 hours using a 3.5 kDa molecular weight cut-off membrane. 250 µL of the dialysed suspension was then added to a 1.5 mL Eppendorf tube to which was added a pre-mixed solution of $CuSO_4$ (100 µM final concentration) and tris(benzyltriazolylmethyl)amine (THPTA, 500 µM final concentration), 5-FAM alkyne (50 µM final concentration), sodium ascorbate (5 mM final concentration) and potassium phosphate buffer (pH 7, 100 mM final concentration to a final reaction volume of 500 µL). Reactions were incubated in a thermoshaker at 30 °C, 1000 rpm (3 mm orbital throw diameter) for 16 hours before dialyzing a 100 µL aliquot against water for 7 days, changing the dialysis water every 24 hours. Samples were diluted to a constant volume of 200 µL and fluorescence measured using a BMGLabtech FluoSTAR Omega plate reader ($\lambda_{ex}$=485 nm, $\lambda_{em}$= 520 nm).

#### 4.5 General procedure for colanic acid extraction

Bacterial cultures were pelleted by centrifugation (70 000 g, 4 °C, 30 minutes) and the resulting supernatant transferred to a clean Falcon tube. 3 volumes of acetone was added to 1 volume of supernatant and incubated at 4 °C overnight. The resulting suspension was centrifuged (13 000 rpm, 4°C, 30 minutes), before carefully removing the supernatant to leave the pellet undisturbed. The pellet was dissolved in Milli-Q $H_2O$ (5-10 mL per 10 mL culture volume) and the resulting solution dialysed against dd$H_2O$ for 16 hours at room temperature, using a 3.5 kDa molecular weight cut-off dialysis membrane. The dialysis cassette was transferred into fresh dd$H_2O$ and dialysed for a further 4 hours at room temperature. The resulting EPS solution was used directly for the colanic acid quantification or freeze-dried to provide solid EPS samples.

### 5 Synthesis

**1,2,3,4-di-*O*-isopropylidene-α-L-galactopyranose**

L-Galactose (245 mg, 1.35 mmol) was suspended in acetone (5 mL) and $ZnCl_2$ (370 mg, 2.7 mmol) was added followed by conc. $H_2SO_4$ (25 µL, 0.46 mmol) and the solution stirred at room temperature for 24 h. To the orange solution was added $NaHCO_3$ (5 mL; sat. aq.) and the solution was stirred for 20 minutes, during which time it went back to colourless. The solution was filtered through celite and the filtrate extracted with ether (3 × 20 mL). The organic phases were combined, dried over $MgSO_4$ and concentrated under reduced pressure to yield the product as a yellow oil which was used without further purification (288 mg, 82% yield); **$R_f$** (3:1 Hexanes/EtOAc) **=** 0.3; **$^1$H NMR** (600 MHZ, $CDCl_3$) δ 5.59 (1H, d, *J* = 5.0 Hz, *H*-1), 4.64 (1H, dd, *J* = 7.9, 2.4 Hz, *H*-3), 4.36 (1H, dd, *J* = 5.0, 2.4 Hz, *H*-2), 4.30 (1H, dd, *J* = 7.9, 1.6 Hz, *H*-4), 3.93 – 3.84 (2H, m, *H*-5, $H$-$6_{\alpha}$), 3.77 (1H, ddd, *J* = 12.5, 6.8, 2.8 Hz, $H$-$6_{\beta}$), 1.56 (3H, s, $CH_3$), 1.48 (3H, s, $CH_3$), 1.36 (6H, br s, 2 × $CH_3$); **$^{13}$C NMR** (126 MHz, $CDCl_3$) δ 109.4 (C), 108.6 (C), 96.3 (CH), 71.6 (CH), 70.8 (CH), 70.6 (CH), 68.0 (CH), 62.3 ($CH_2$), 26.0 ($CH_3$), 25.9 ($CH_3$), 24.9 ($CH_3$), 24.3 ($CH_3$). Spectroscopic data matches the literature[6].

**6-azido-1,2,3,4-di-*O*-isopropylidene-6-deoxy-α-L-galactopyranoside**

To 1,2,3,4-di-*O*-isopropylidene-α-L-galactopyranose (288 mg, 1.1 mmol) was added anhydrous DCM (5 mL) and DIPEA (1.3 mL, 7.5 mmol) under argon and the solution cooled on ice. $Tf_2O$ (0.5 mL, 3.0 mmol) was added to the solution dropwise over 5 minutes and the solution stirred for a further 15 minutes. TLC (3:1 Hexanes/EtOAc, visualised using permanganate) showed product ($R_f$ = 0.8) and remaining starting material ($R_f$ = 0.3). A further portion of $Tf_2O$ (0.25 mL, 1.5 mmol) was added dropwise to the solution, whereupon TLC confirmed complete consumption of starting material. Cold $H_2O$ (20 mL) was added to the solution which was extracted with DCM (3 × 20 mL). The organic phases were combined, dried over $Na_2SO_4$ and concentrated under reduced pressure to yield crude 1,2:3,4-

di-*O*-isopropylidene-6-*O*-(trifluoromethanesulfonyl)-α-L-galactopyranose as a red oil. The oil was dissolved in DMSO (5 mL), $NaN_3$ (0.7 g, 10.8 mmol) was added and the solution stirred at room temperature for 20 hours. $H_2O$ (30 mL) was added and the solution was extracted with $Et_2O$ (3 × 20 mL). The combined organic phases were dried over $MgSO_4$ and concentrated under reduced pressure. Purification by flash chromatography using 9:1 Hexanes/EtOAc gave the product as a clear oil (200 mg, 64% yield); **$R_f$** (9:1 Hexanes/EtOAc) = 0.2; **$^1$H NMR** (500 MHz, $CDCl_3$) δ 5.57 (1H, d, *J* = 5.0 Hz, *H*-1), 4.65 (1H, dd, *J* = 7.9, 2.5 Hz, *H*-3), 4.36 (1H, dd, *J* = 5.0, 2.5 Hz, *H*-2), 4.22 (1H, dd, *J* = 7.9, 2.0 Hz, *H*-4), 3.94 (1H, ddd, *J* = 7.5, 5.3, 2.0 Hz, *H*-5), 3.53 (1H, dd, *J* = 12.7, 7.9 Hz, $H$-6$_a$), 3.39 (1H, dd, *J* = 12.7, 5.3 Hz, $H$-6$_b$), 1.57 (3H, s, $CH_3$), 1.48 (3H, s, $CH_3$), 1.37 (3H, s. $CH_3$), 1.36 (3H, s, $CH_3$); **$^{13}$C NMR** (126 MHz, $CDCl_3$) δ 109.6 (C), 108.8 (C), 96.4 (CH), 71.2 (CH), 70.8 (CH), 70.4 (CH), 67.0 (CH), 50.7 ($CH_2$), 26.0 ($CH_3$), 26.0 ($CH_3$), 24.9 ($CH_3$), 24.44 ($CH_3$). Spectroscopic data matches the literature[7].

**6-Azido-L-fucose**

6-azido-1,2,3,4-di-*O*-isopropylidene-6-deoxy-α-L-galactopyranoside (200 mg, 0.7 mmol) was dissolved in TFA (5 mL; 80% aq.) and stirred at room temperature for 3 hours. The solution was concentrated under reduced pressure yielding a yellow oil. The product was obtained by recrystallisation from EtOH (1 mL) as a white solid. The recrystallisation liquor was concentrated to ~200 µL and further product was isolated by layer diffusion crystallisation with ether (84 mg, 58% yield); **$R_f$** (15% MeOH in DCM) = 0.2; **$\alpha_D$** = -66.4 ° (*c* = 0.8, $H_2O$), [Lit[8] = -56.4 °, (*c* = 1, $H_2O$)]; **$^1$H NMR** (600 MHz, $D_2O$) δ 5.30 (1H, dd, *J* = 8.2, 3.8 Hz, α*H*-1), 4.64 (1H, d, *J*=7.9 Hz, β*H*-1), 4.23 (1H, ddd, *J* = 8.9, 4.4, 1.1 Hz, α*H*-5), 3.99 (1H, dd, *J* = 3.3, 1.2 Hz, α*H*-4), 3.93 (1H, dd, *J* = 3.4, 1.1 Hz, β*H*-4), 3.89 (1H, dd, *J* = 10.3, 3.4 Hz), 3.85 (1H, ddd, *J*=8.6, 4.4, 1.1 Hz, β*H*-5), 3.82 (1H, dd, *J* = 10.3, 3.6 Hz, α*H*-2), 3.68 (1H, dd, *J* = 9.9, 3.5 Hz, β*H*-3), 3.62 (1H, dd, *J*=13.0, 8.5 Hz, β$H$-6$_a$), 3.57 (1H, dd, *J* = 13.0, 8.6 Hz, α$H$-6$_a$), 3.54 – 3.48 (3H, m, β*H*-2, α$H$-6$_b$ + β$H$-6$_b$); **$^{13}$C NMR** (126 MHz, $D_2O$) δ 96.4 (CH-β), 92.4 (CH-α), 73.4 (CH-β), 72.66 (CH-β), 71.7 (CH-β), 69.6 (CH-α), 69.1 (CH-β), 69.0 (CH-α), 68.9 (CH-α), 68.2 (CH-α), 50.9 (CH-α), 50.7 (CH-β); **HRMS** (ESI+) $[M+H]^+$ found 206.0782, $C_6H_{12}N_3O_5$ requires 206.0771. Spectroscopic data matches the literature[9].

#### 6 Supplementary data

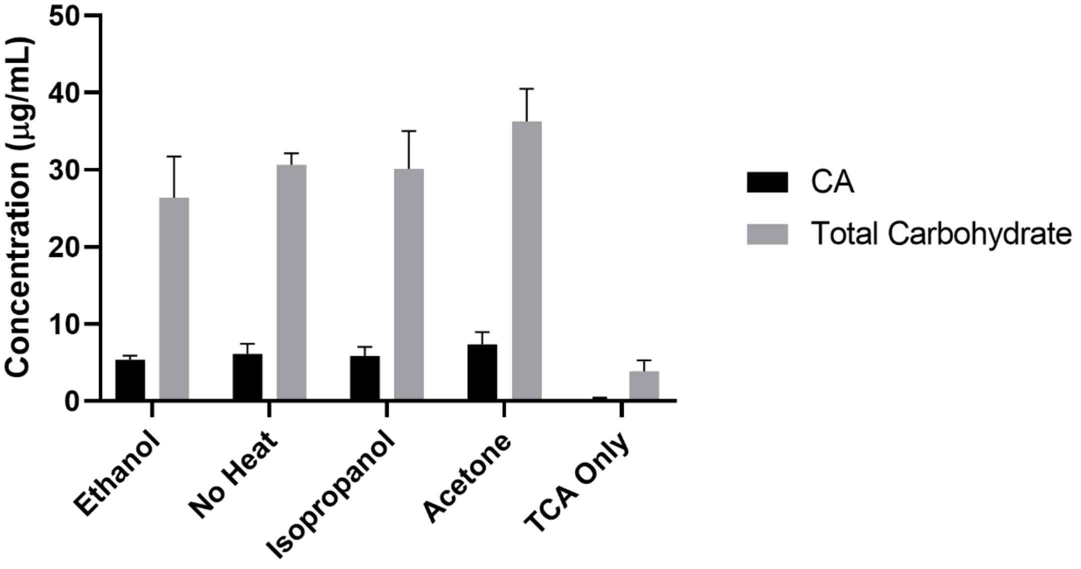

***Figure S5.*** *Effect of the extraction procedure and solvent used in the CA production (black) and the total carbohydrate content (grey). Error bars represent the standard deviation of values from three independent biological replicates. TCA: trichloroacetic acid.*

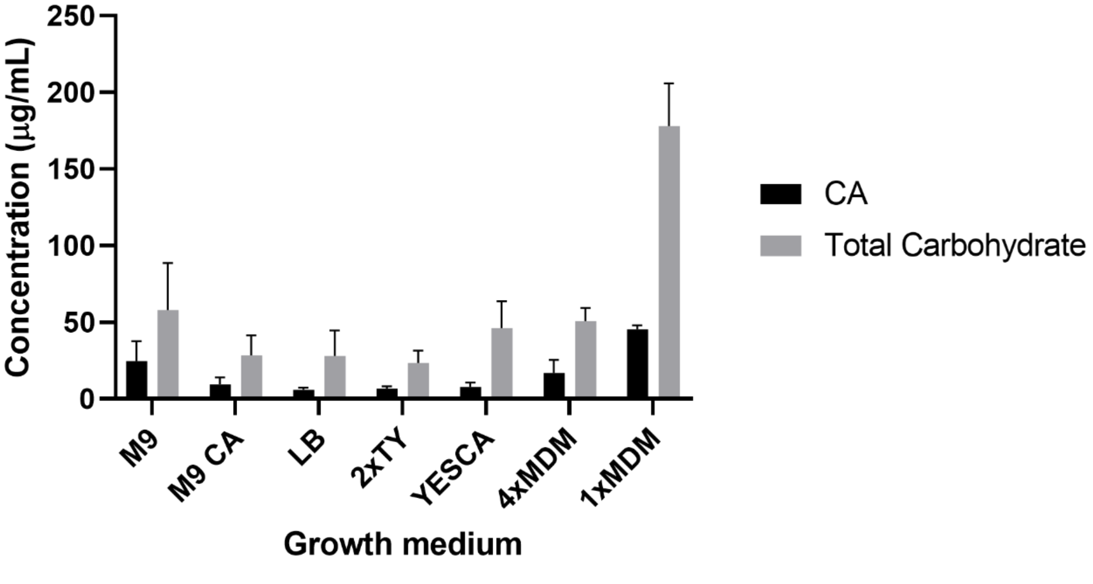

***Figure S6.*** *Effect of the media used for the growth of E. coli JM109_pRcsA in the production of CA (black) and total carbohydrate content (grey). Error bars represent the standard deviation of values from three independent biological replicates.*

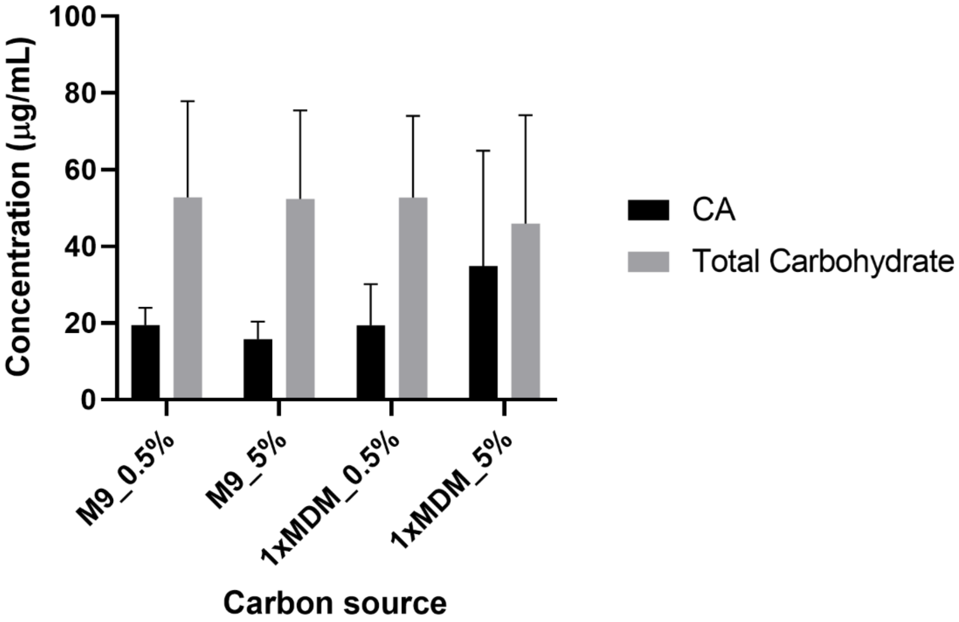

***Figure S7.*** *Effect of the glucose percentage used either in M9 or 1xMDM (0.5% or 5%) in the production of CA (black) and total carbohydrate production (grey). Error bars represent the standard deviation of values from three independent biological replicates.*

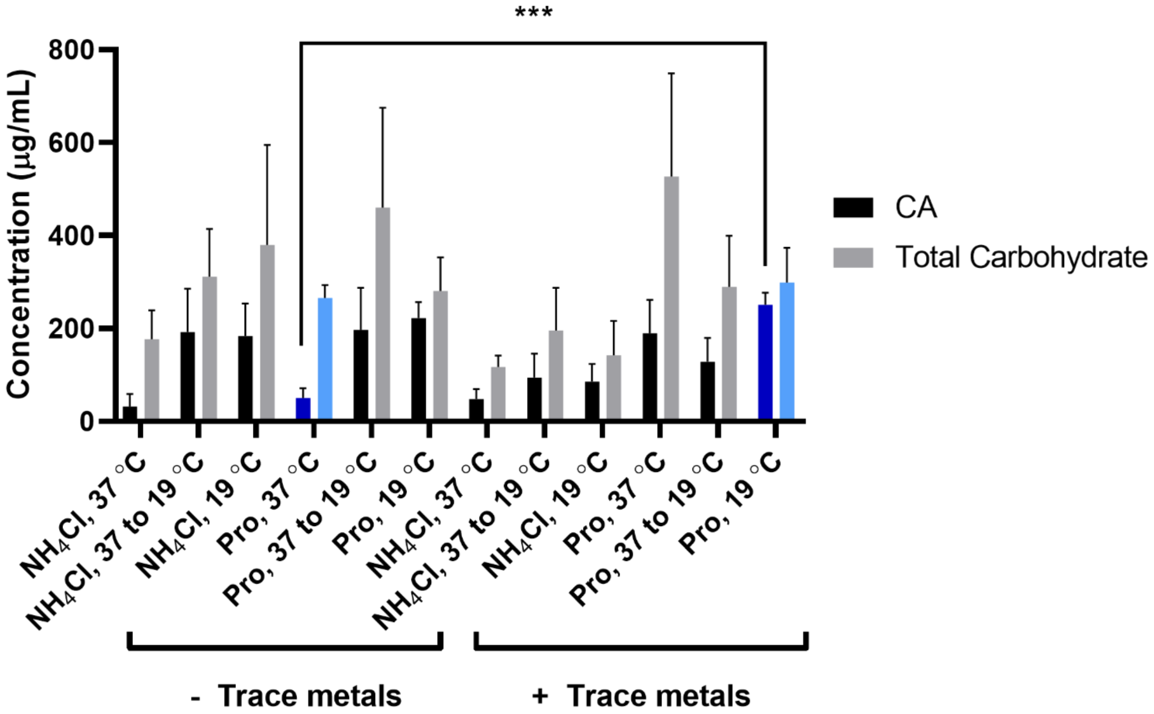

***Figure S8.*** *Effect of the incubation temperature and the addition of Cu(II) and Fe(II) as trace metal ions in the production of CA (black) and the total carbohydrate content (grey). Data from the proline control cultures and proline cultures incubated at 19 °C with the addition of metal ions are highlighted in blue. Error bars represent the standard deviation of values from three independent biological replicates; ***P<0.0001 (Dunnet's test).*

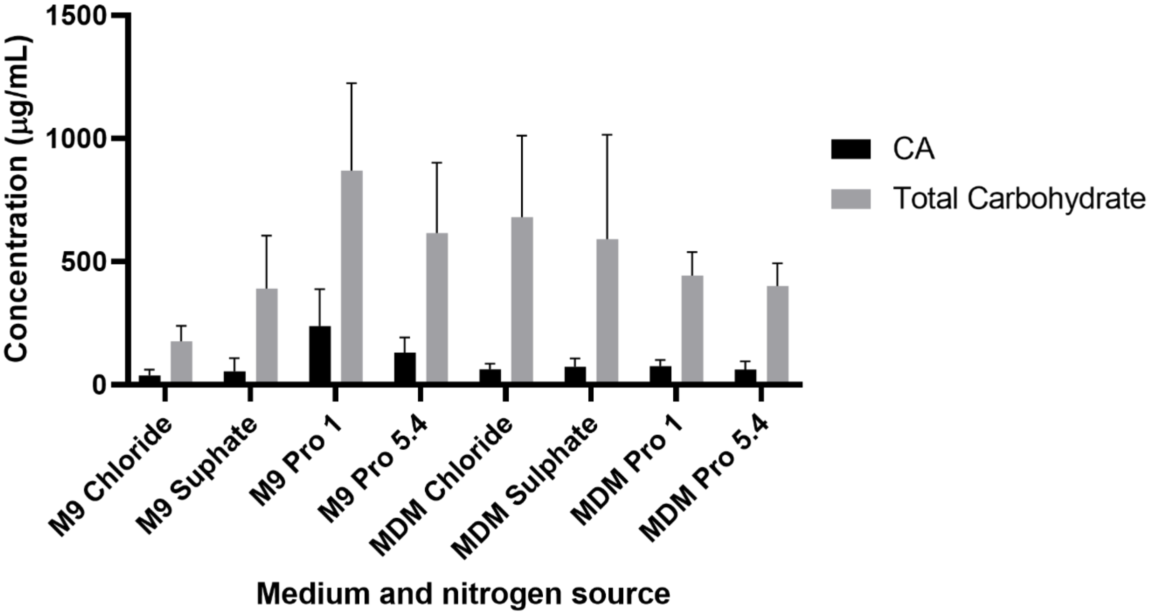

***Figure S9.*** *Effect of the nitrogen source used in the growth media in the production of CA (black) and the total carbohydrate content (grey). Error bars represent the standard deviation of values from three independent biological replicates.*

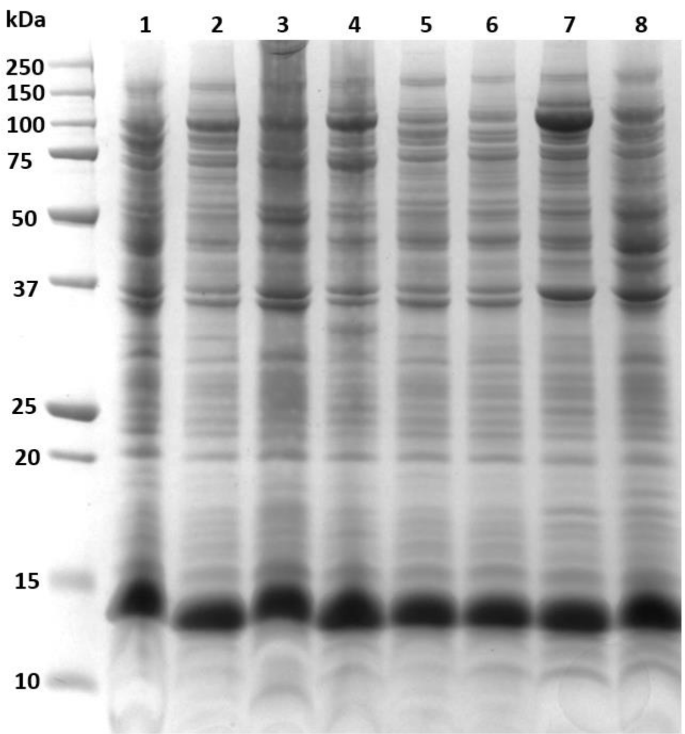

| Lane | Strain | Pre-induction temp. (°C) | Media |
|---|---|---|---|
| **1** | JM109(DE3)Δ*gmd-fcl*_pRcsA_pFkp | 22 | M9-EPS |
| **2** | JM109(DE3)Δ*gmd-fcl*_pRcsA_pFkp | 37 | M9-EPS |
| **3** | JM109(DE3)Δ*gmd-fcl*_pRcsA_pFkp | 22 | LB |
| **4** | JM109(DE3)Δ*gmd-fcl*_pRcsA_pFkp | 37 | LB |
| **5** | JM109(DE3)Δ*gmd-fcl*_pRsfDuet1 | 37 | M9-EPS |
| **6** | JM109(DE3)Δ*gmd-fcl*_pRsfDuet1 | 37 | LB |
| **7** | BL21(DE3)_pFkp | 37 | LB |
| **8** | BL21(DE3)_pFkp | 22 | LB |

***Figure S10.*** *SDS-PAGE analysis of Fkp expression. All cultures were incubated at 22 °C (room temperature) for 24 hours post-induction with IPTG. JM109(DE3)Δgmd-fcl_pRsfDuet1 was included as a negative control for each condition; BL21(DE3)_pFkp was included as a positive control for each condition. Theoretical size of Fkp: 106 kDa.*
